# Islet amyloid polypeptide tagged with green fluorescent protein localises to mitochondria and forms filamentous aggregates in *Caenorhabditis elegans*

**DOI:** 10.1101/2023.10.27.564377

**Authors:** Mehmet Akdag, Vera van Schijndel, Tessa Sinnige

## Abstract

Type 2 diabetes (T2D) is the most common form of diabetes and represents a growing health concern. A characteristic feature of T2D is the aggregation of islet amyloid polypeptide (IAPP), which is thought to be associated with the death of pancreatic β-cells. Inhibiting IAPP aggregation is a promising therapeutic avenue to treat T2D, but the mechanisms of aggregation and toxicity are not yet fully understood. *Caenorhabditis elegans* is a well-characterised multicellular model organism that has been extensively used to study protein aggregation diseases. In this study, we aimed to develop a simple *in vivo* model to investigate IAPP aggregation and toxicity based on expression in the *C. elegans* body wall muscle cells. We show that IAPP tagged with green fluorescent protein (GFP) localises to mitochondria in this tissue, in line with previous observations in mouse and human pancreatic β-cells. The IAPP-GFP fusion protein forms solid aggregates, which have a filamentous appearance as seen by electron microscopy. However, the animals do not display a strong motility phenotype, suggesting that the IAPP-GFP aggregates are not considerably toxic. Nevertheless, the mitochondrial localisation and aggregate formation may be useful read-outs to screen for IAPP-solubilizing compounds as a therapeutic strategy for T2D.

**Graphical abstract:** 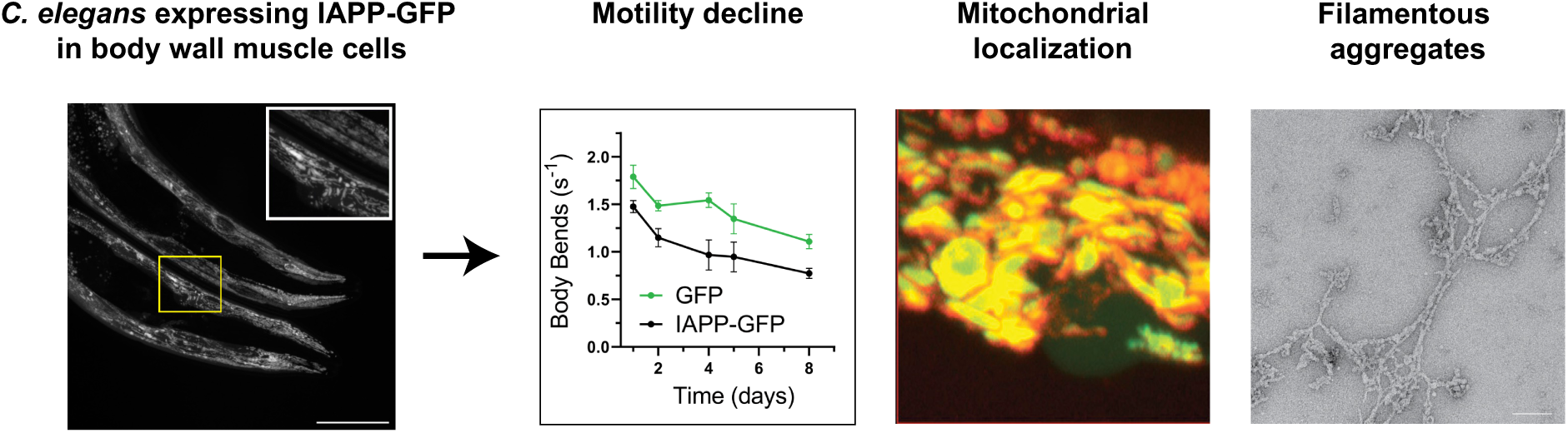

## Introduction

Diabetes is affecting the lives of about 537 million people globally and was associated with 6.7 million deaths in 2021 [1]. The most prevalent form of diabetes is Type 2 diabetes (T2D) in which patients become hyperglycaemic and insensitive to insulin, causing increased levels of counteracting insulin and the co-secreted islet amyloid polypeptide (IAPP), or amylin, that together regulate glucose homeostasis [2].

IAPP is produced as an 89 amino acid long preprohormone (preproIAPP). Upon reaching the secretory pathway, the signal peptide is cleaved off forming proIAPP. Prior to secretion, the flanking regions of proIAPP are cleaved, the peptide is amidated at the C-terminus and an intramolecular disulfide bond is formed to yield the 37 amino acid long active mature IAPP hormone [3]. IAPP is a highly amyloidogenic peptide that leads to the formation of amyloid deposits in around 90 % of T2D patients. These IAPP aggregates are associated with the death of insulin secreting pancreatic β-cells, which is a major histopathological characteristic of T2D [4–6].

The mechanisms of IAPP aggregation have previously been studied using *in vitro* approaches. IAPP aggregation follows a nucleation-dependent pathway similar to other amyloidogenic proteins. *In vitro* kinetics assays revealed that the driving mechanism for IAPP fibril formation is secondary nucleation, a process in which new fibrils are formed by catalytic events on already existing fibrils [7–9]. IAPP can also interact with membranes and its aggregation disturbs lipid bilayers, which is thought to contribute to its toxicity [10–12]. However, it is unclear to what extent these *in vitro* studies translate to the complex environment *in vivo*. Although mouse models are available to study diabetes [13,14], there is a need for a more basic tool to address the mechanisms of IAPP aggregation and toxicity, and to use for drug screening.

The nematode *Caenorhabditis elegans* is a well-characterised multicellular model organism that has been extensively used to study protein aggregation, neurodegeneration and ageing. *C. elegans* models based on the transgenic expression of human amyloidogenic proteins such as amyloid-β (Aβ), α-synuclein, and polyQ repeats have been widely used in the field [15–19]. Expression of these proteins in the body wall muscle cells is associated with toxicity as judged from impaired motility of the animals, which has been used as the basis for genetic and drug screening [20–22]. The muscle cells are furthermore relatively large, allowing for straight-forward visualisation of protein aggregates by fluorescence microscopy [18,19,23].

IAPP aggregation has been less well studied in *C. elegans* in comparison, with only a few models reported in the literature. It was shown that heat-shock inducible expression of mature IAPP led to changes in the transcriptome in addition to developmental defects [24]. Expression of preproIAPP tagged with yellow fluorescent protein (YFP) in various tissues resulted in aggregates visible by microscopy and was also associated with a developmental phenotype [25]. However, a simple model system in which human mature IAPP is expressed in the body wall muscle cells is not currently available, and here we set out to develop and characterise such a model.

## Results

In order to develop a *C. elegans* model to study IAPP aggregation and toxicity, we designed a construct encoding human mature IAPP flanked by green fluorescent protein (GFP) separated by a 9 amino acid linker sequence. We used the *unc-54* promoter to express the IAPP-GFP fusion protein specifically in the body wall muscle cells, and we created two genomic integrants. As a control strain, we also generated a body wall muscle cell specific cytosolic GFP expressing strain (**Figure 1A**). We confirmed IAPP-GFP expression in the body wall muscle cells by microscopy (**Figure 1B**) and by western blot with an anti-GFP antibody (**Figure S1**). We did not observe a significant difference in protein levels between the two IAPP-GFP strains.

**Figure 1:**
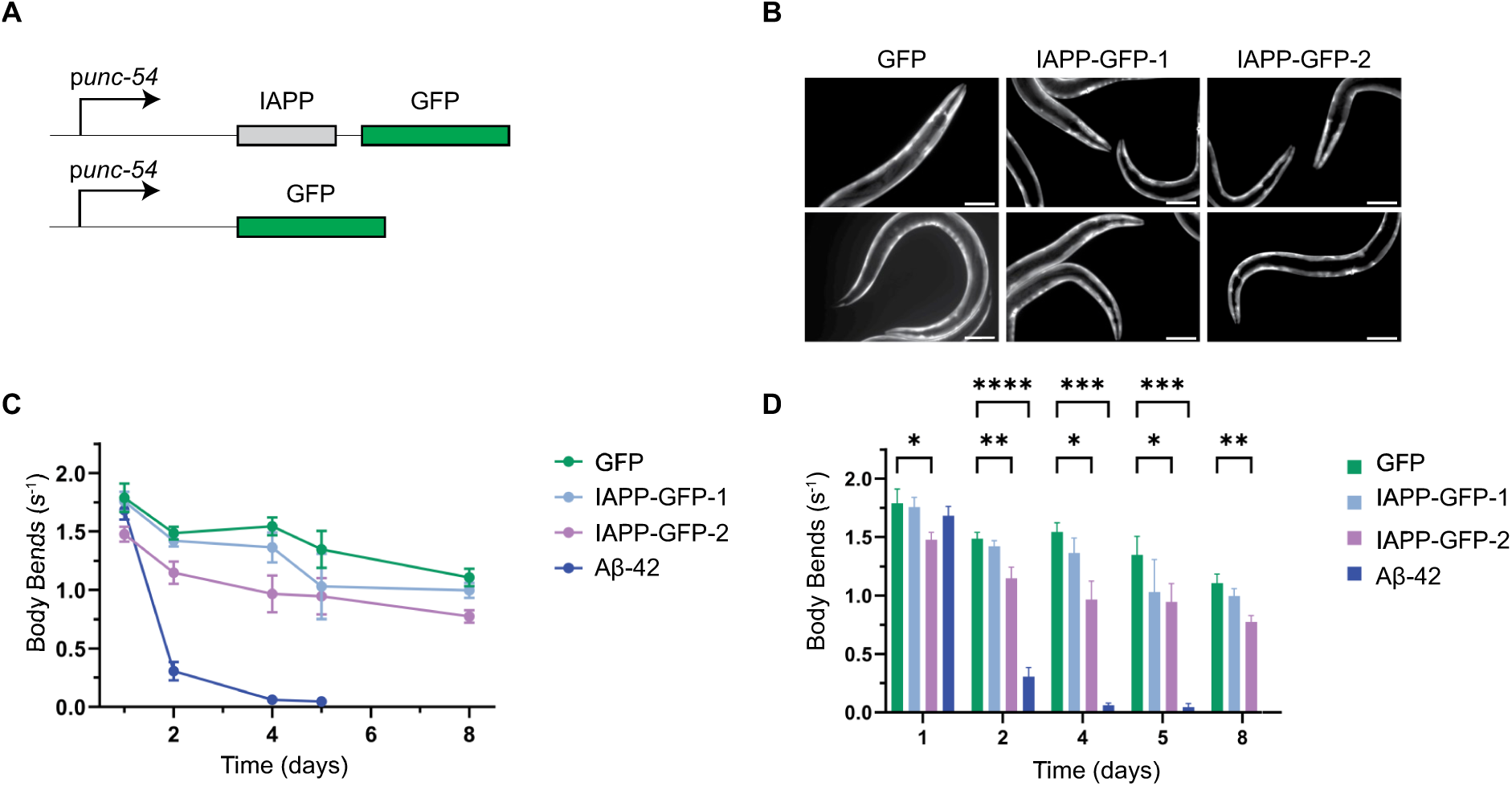
Developing a body wall muscle cell specific *C. elegans* IAPP-GFP model. **(A)** Schematic design for IAPP-GFP and GFP expression constructs. **(B)** Fluorescent microscopy images confirming expression of IAPP-GFP and GFP in the body wall muscle cells. (Scale bar 100 µm) **(C)** Motility assay of the IAPP-GFP strains, GFP and Aβ-42 controls. Data collection and analysis were done in a blinded manner and at least 30 animals were used for each data point. **(D)** Quantification of the motility assay. Two-way ANOVA analysis with Dunnet multiple comparison test was employed for statistical analysis. (* p < 0.05; ** p < 0.01)

We proceeded to use motility assays to assess the toxicity of IAPP-GFP expression. Nematodes expressing Aβ-42 in body wall muscle cells and wild-type (N2) animals were used as controls. Thrashing in liquid medium was recorded at different time points from day 1 up to day 8 of adulthood. The animals were incubated at 25 °C throughout adulthood to promote protein aggregation and toxicity, and indeed Aβ-42 expressing worms showed severe motility defects at this temperature consistent with previous reports (**Figure 1C, D**) [16]. Compared to the GFP control strain, both IAPP-GFP strains performed somewhat worse, but only one IAPP-GFP integrant displayed a significant reduction in motility **(Figure 1C, D).** The Aβ-42 strain performs considerably worse, indicating that IAPP-GFP expression is not as toxic.

To test whether the relatively large fluorophore tag may interfere with the properties and toxicity of IAPP, we also generated strains that express untagged IAPP. We used different signal sequences to target the peptide to the secretory pathway and inserted an SL2 splicing site between IAPP and GFP. This way the mRNA encoding IAPP-SL2-GFP is spliced and both proteins are translated separately (**Figure S2A**). Although we confirmed expression based on the presence of GFP in these strains (**Figure S2B**), we did not detect any toxicity with motility assays **(Figure S2C, S2D)**. We concluded that either untagged IAPP does not cause any toxicity in the body wall muscle cells, or the levels are too low to elicit observable effects. Therefore we proceeded our study with the IAPP-GFP strains.

We further characterised the IAPP-GFP strains using confocal microscopy to find out if the mild motility defect could be caused by protein aggregation. While the GFP control showed diffuse expression throughout the muscle cells (**Figure 2A**), both of the IAPP-GFP strains showed a heterogeneous distribution forming elongated and punctate structures (**Figure 2B, 2C**). We hypothesised that this pattern may correspond to either solid or liquid-like assemblies, or alternatively they could arise from localisation to certain organelles.

**Figure 2:**
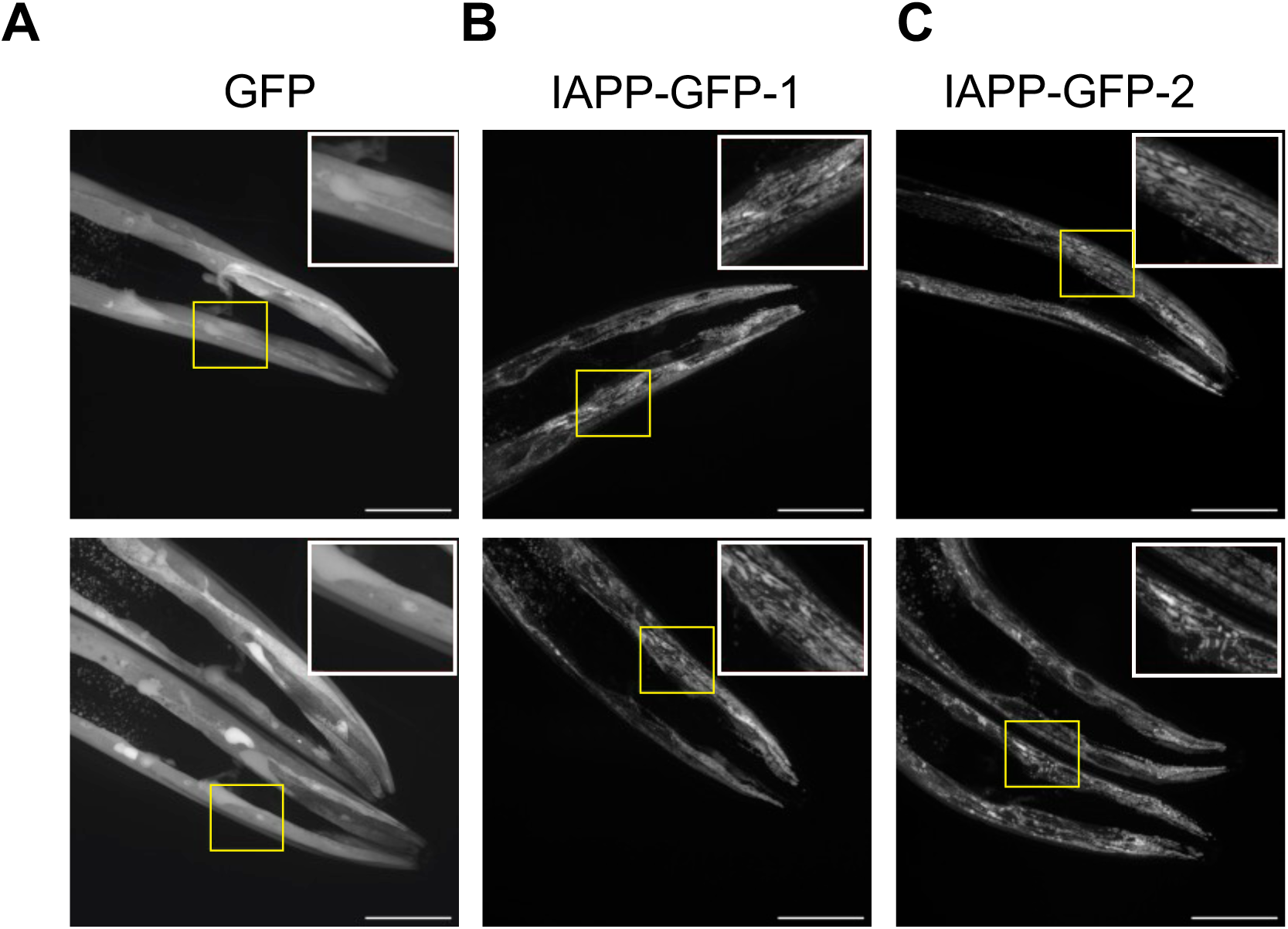
IAPP-GFP strains show a distinct heterogeneous distribution as seen by confocal microscopy. Example images of the *C. elegans* head regions are shown for **(A)** GFP, **(B)** IAPP-GFP-1, **(C)** IAPP-GFP-2. Yellow boxes are enlarged at the top right of each image. Animals were imaged on day 5 of adulthood. (Scale bar 50 µm)

Previously, IAPP was found to associate with mitochondria in mouse pancreatic islets and in human insulinoma cells [26,27]. To find out whether the distribution of IAPP-GFP in our *C. elegans* model can be explained by co-localisation with mitochondria, we crossed IAPP-GFP-2 worms with a mitochondrial marker strain that expresses TOMM20-mKate2 in the body wall muscle cells [28]. In the control strain, we did not observe co-localisation of cytosolic GFP and TOMM20. On the other hand, the IAPP-GFP signal co-localised at least partially with the mitochondrial membrane in the body wall muscle cells (**Figure 3**). It should be noted that the expression levels of TOMM20-mKate2 are relatively low, which may explain the apparent lack of co-localisation in some of the head muscle cells.

**Figure 3:**
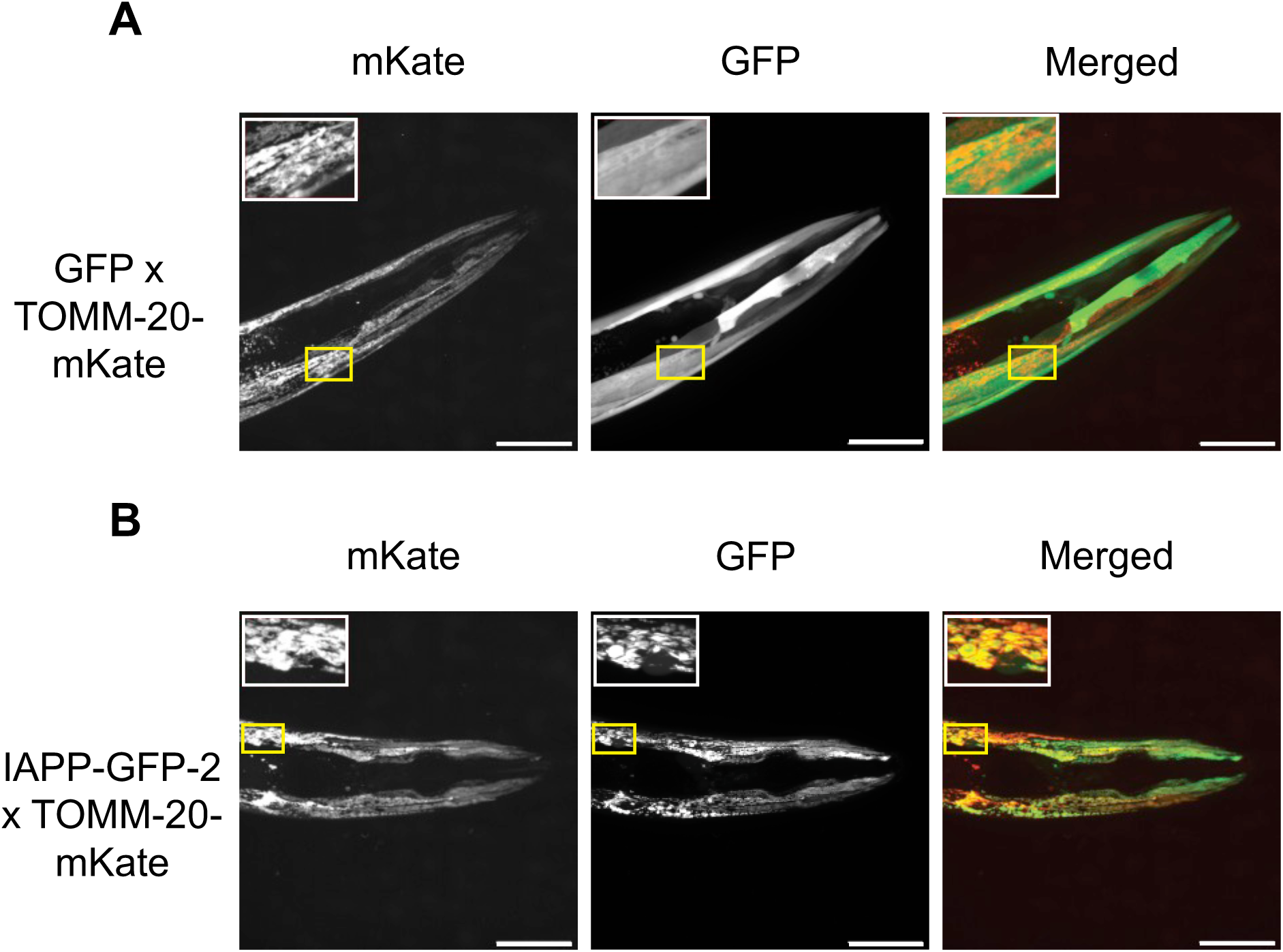
Confocal imaging shows partial IAPP-GFP co-localisation to the mitochondrial membrane. **(A)** GFP and **(B)** IAPP-GFP-2 animals were crossed with the TOMM-20-mKate mitochondrial membrane marker expressing strain and the offspring were imaged at day 2 of adulthood. (Scale bar 50 µm)

We then investigated if IAPP-GFP forms solid aggregates using fluorescence recovery after photobleaching (FRAP). While the cytosolic GFP signal instantly recovers after photobleaching, both IAPP-GFP strains showed only limited recovery indicative of solid aggregates (**Figure 4A, 4B**). We then validated the presence of insoluble IAPP-GFP aggregates using a fractionation assay. Whole lysate from IAPP-GFP animals was separated into SDS soluble and insoluble fractions. We observed that IAPP-GFP was partially recovered in the SDS insoluble fraction (**Figure 4C**), while GFP remained in the soluble fraction. Furthermore, treatment with 1,6-hexanediol, which is commonly used to disturb liquid condensates, did not alter the distribution of IAPP-GFP (**Figure S3**). Altogether these data demonstrate that IAPP-GFP forms solid aggregates when expressed in *C. elegans* body wall muscle cells.

**Figure 4:**
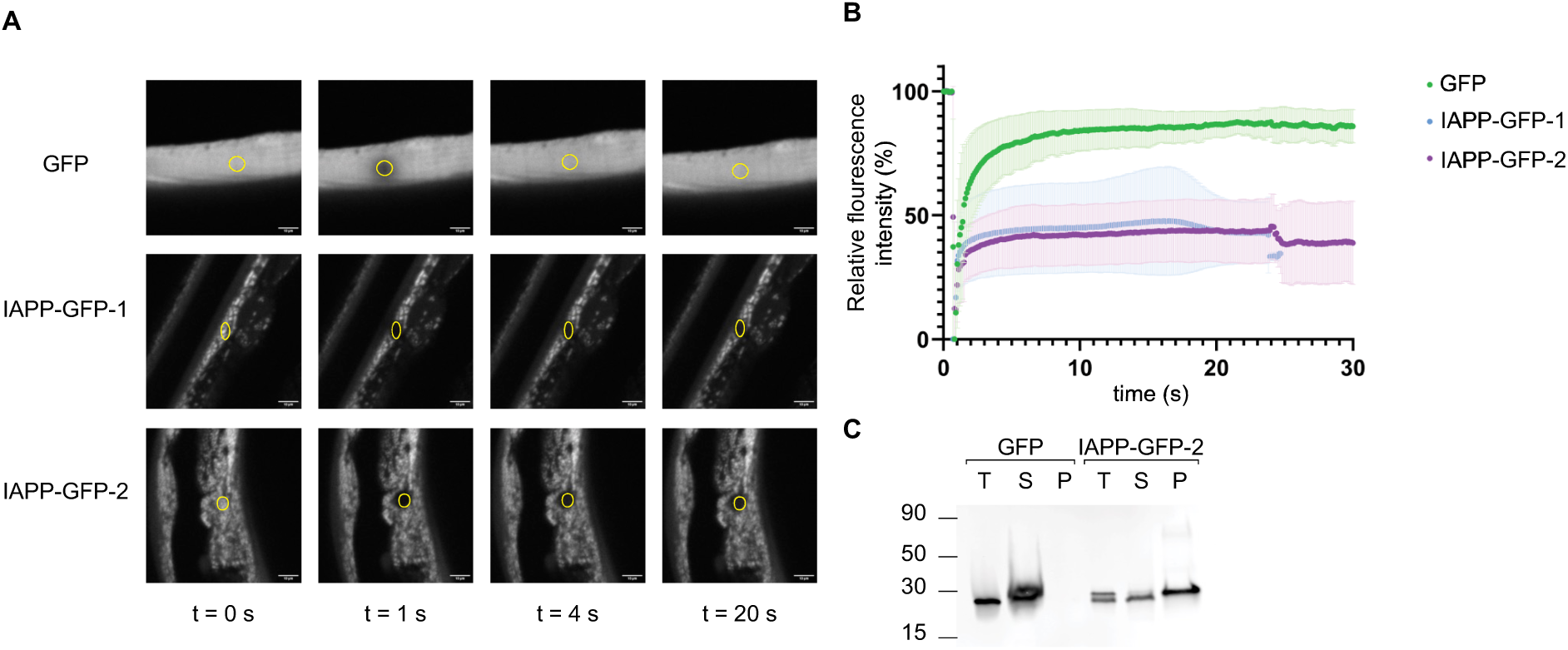
IAPP-GFP forms solid aggregates as observed by microscopy and biochemical assays. **(A)** FRAP of GFP (top), IAPP-GFP-1 (middle) and IAPP-GFP-2 (bottom). Yellow circles indicate the photobleached region. (Scale bar 10 µm) **(B)** Quantification of fluorescence recovery was achieved by measuring signal intensities in the photobleached areas. (N = 4-8 animals per strain, data presented as the mean± standard deviation) **(C)** Western blot of fractionated lysates probed with an anti-GFP antibody. Total lysates (T), supernatant (S) and pellet (P) are shown. Only IAPP-GFP accumulates in the SDS-insoluble fraction (P).

We further investigated the nature of the IAPP-GFP aggregates by staining with the amyloid-binding dye NIAD-4 [29] (**Figure S4**). We used Aβ-42 animals as a positive control and observed deposits stained with NIAD-4. However, in the IAPP-GFP strains, we did not observe any NIAD-4 signal, suggesting that the aggregates may be amorphous in nature. Alternatively, they might be in a fibrillar form that is not recognised by NIAD-4.

In order to distinguish between these possibilities, we extracted IAPP-GFP by immunoprecipitation and analysed the morphology using negative staining transmission electron microscopy (TEM) **(Figure 5)**. In line with our previous FRAP and fractionation results, electron microscopy verified that IAPP-GFP forms aggregates in *C. elegans*. TEM micrographs of IAPP-GFP aggregates showed that they form networks of filamentous structures (**Figure 5A, B**). The filaments are variable in width and length, partially decorated with bead-like structures, and interspersed with less ordered structures. This irregular morphology may explain why NIAD-4 does not recognise the IAPP-GFP aggregates, and perhaps also their limited toxicity. In addition to the filamentous structures, we also observed spherical particles with diameters of around 15-25 nm which likely correspond to soluble aggregate species (**Figure 5C**). None of these structures were found in the GFP control sample (not shown).

**Figure 5:**
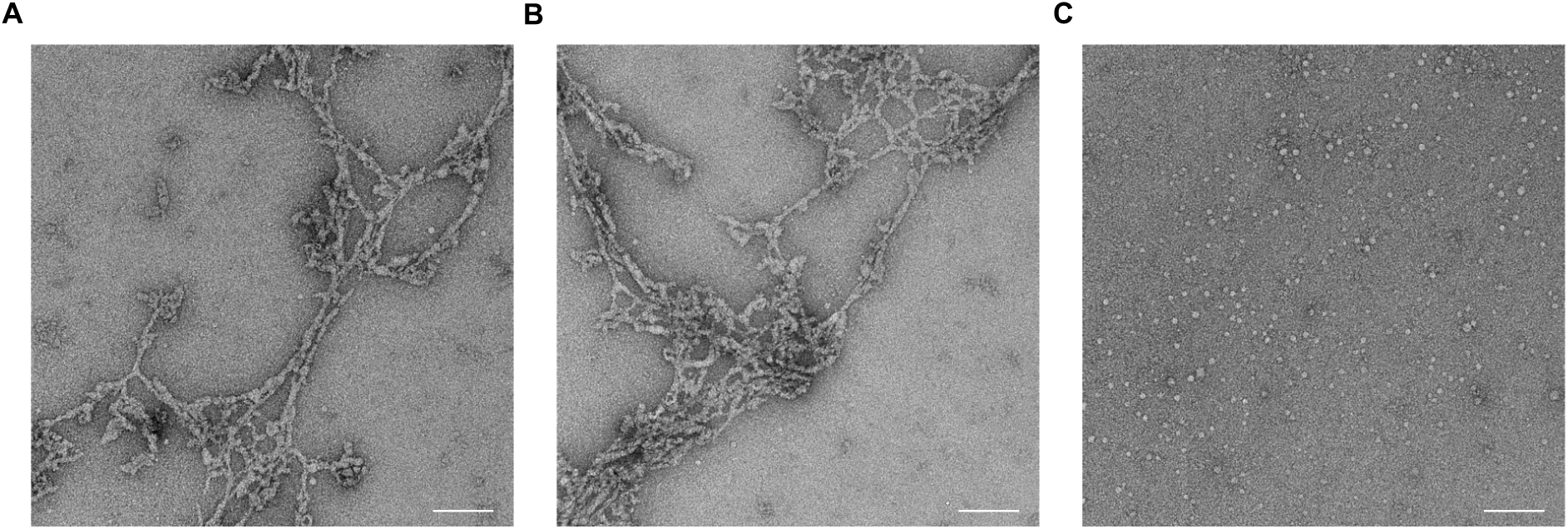
Transmission electron micrographs of negatively stained samples of IAPP-GFP purified by immunoprecipitation. IAPP-GFP can be found to form networks of filamentous structures **(A, B)** as well as spherical particles **(C)**. (Scale bar 200 nm)

## Discussion

In this study, we presented a novel *C. elegans* model expressing GFP-tagged IAPP in the body wall muscle cells. IAPP-GFP appears to spontaneously associate with mitochondria, in the absence of a targeting sequence (**Figure 3**). These observations are in line with our recent *in vitro* results showing that IAPP strongly interacts with anionic lipids including cardiolipin, which is abundant in the mitochondrial membrane [12]. It has also been suggested that the negative curvature induced by phosphatidylethanolamine and cardiolipin promotes IAPP binding to the mitochondrial membrane [26]. IAPP was found to be associated with mitochondria in mouse pancreatic β-cells and human insulinoma cells [26,27], and IAPP aggregates positive for the A11 antibody were found to compromise mitochondrial integrity [27]. From *in vitro* studies, it has also become clear that IAPP aggregation, and in particular fibril elongation, causes membrane disturbances [8,10,12]. However, it is unlikely that IAPP-GFP aggregation causes considerable damage to the mitochondrial membrane in the muscle cells of our *C. elegans* model, as this would be expected to induce strong motility defects.

Next to the large filamentous aggregates, we also observed spherical particles in the IAPP-GFP sample by TEM (**Figure 5**). Interestingly, IAPP has been previously shown to form micelles with diameters of 12-15 nm [30]. GFP is about 3 nm in diameter, so it is conceivable that the particles we observe correspond to IAPP micelles decorated with GFP. However, these are apparently not considerably toxic either. We observed that IAPP-GFP consistently ran as a double band on western blots (**Figure S1, Figure 4C**), the lower band corresponding to the molecular weight of GFP suggestive of a cleavage product. While we cannot exclude that this cleavage took place during preparation of the lysate, it is possible that a mixture of tagged and free IAPP is present in our *C. elegans* model. Still, the motility phenotype is surprisingly mild.

Both of the previously reported *C. elegans* IAPP models were found to have a developmental defect, which we did not observe in our models (not shown). In one study, preproIAPP was expressed as a YFP fusion protein [25], suggesting that the presence of the tag does not necessarily interfere with toxicity, but that the longer construct may be more detrimental. In the other study, the mature peptide was expressed without a tag under the control of a heat shock promoter, which drives expression in neurons, pharyngeal muscle, and the hypodermis upon heat induction [24]. Both the expression pattern and the heat stress may have contributed to the observed developmental delay in this study.

Although *C. elegans* has been widely used in the protein aggregation field, the physiology of the nematode is limited in complexity and does not allow human diseases to be fully recapitulated. With regards to T2D, *C. elegans* does not have a pancreas and the human machinery to secrete and process preproIAPP is not entirely conserved. *C. elegans* has other tissues capable of secretion including the neurons that secrete the insulin homolog. Whereas the purpose of the current study was to establish a simple model based on IAPP expression in the body wall muscle cells, expression in the neurons may be an interesting avenue to develop a more physiological model for the aggregation of secreted IAPP, in particular in combination with sub-stoichiometric labelling as recently demonstrated for Aβ [31]. In fact, expression of tagged preproIAPP in the *C. elegans* neurons led to its secretion and uptake in the macrophage-like coelomocyte cells as shown in one of the previous studies [25]. However, it is not clear to what extent this protein construct was processed, given that removal of the C-terminal part of proIAPP would result in loss of the fluorescent tag. As such, it remains a challenge to develop a faithful *C. elegans* model to investigate IAPP aggregation and toxicity.

In conclusion, our study shows that an IAPP-GFP fusion protein forms solid, filamentous aggregates in *C. elegans* body wall muscle cells, which appear to co-localise with mitochondria. Although the animals display only a modest motility phenotype, the presence of solid aggregates and mitochondrial localisation may be useful as read-outs to test IAPP solubilizing drugs in future studies.

## Methods

### Plasmid generation

DNA plasmids were generated using Gibson Assembly (NEB). The *unc-54* promoter and *unc-54* 3’-UTR sequences were amplified from *C. elegans* genomic DNA. DNA fragments encoding human mature IAPP along with signal sequences (ss-IAPP, ins1-IAPP and mature IAPP) were synthesised by IDT. The SL2 trans-splicing sequence was amplified from plasmid pTS06 [32]. The linker sequence for the fusion protein was incorporated in the primer used to amplify the GFP sequence by PCR. Inserts were assembled into the pDEST plasmid backbone using the Gibson Assembly manufacturer’s protocol. All constructs were verified by Sanger sequencing. Primer sequences and plasmids used in this study are given in the supplementary information (**Table S1**, **S2** and **S3)**.

### *C. elegans* strains and maintenance

Nematodes were grown on nematode growth media (NGM) plates seeded with *Escherichia coli* OP50 at 20 °C, unless stated otherwise.

Transgenic animals were generated by microinjections. 100 ng/µL injection mix (70 ng/µL 1 kb DNA ladder (NEB), 30 ng/µL plasmid of interest) was injected into the gonads of adult hermaphrodite wild-type N2 animals. To integrate the transgene into the genome, GFP-expressing L4-staged animals were treated with 30 mJ/cm^2^ UV light using a UVP Crosslinker CL-3000 (Analytik Jena). UV-treated animals were incubated at 20 °C for 2 weeks and fluorescently labeled offspring were singled and screened to obtain integrant lines. Integrant lines were backcrossed with N2 five times to eliminate any UV-induced mutations.

Age synchronisation was achieved by incubating gravid adults on *E. coli* OP50-seeded NGM plates at 20 °C for 1 h. Adults were removed and eggs were incubated at 20 °C until they reached adulthood.

### *C. elegans* strains used in this study: N2 (Bristol)

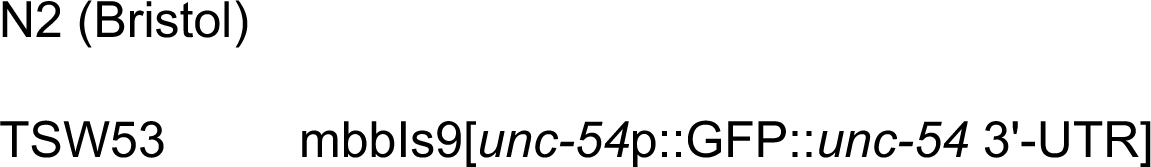
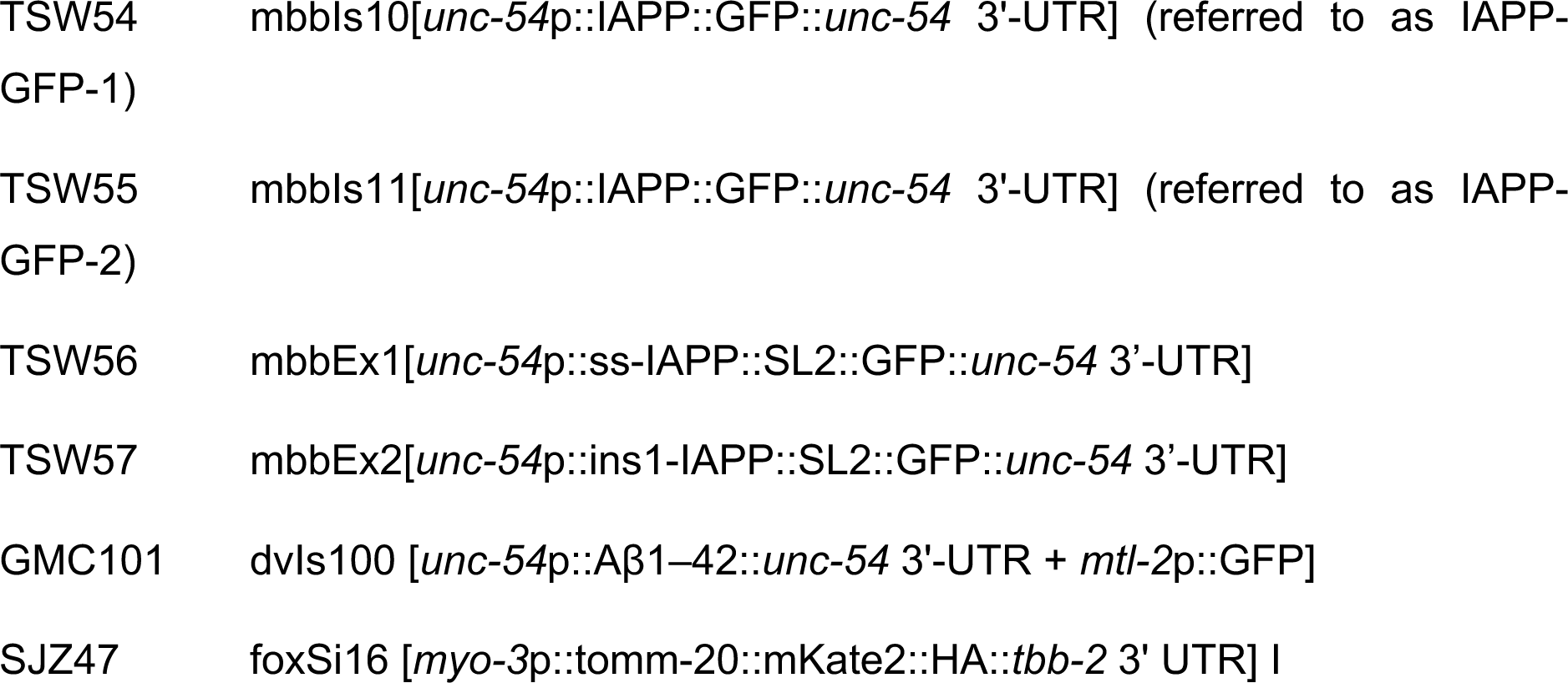

### Thrashing assay

For motility assays, age-synchronised nematodes were grown at 20 °C until they reached adulthood day 1, then the temperature was switched to 25 °C to promote aggregation and toxicity. Animals were transferred to M9 buffer (22 mM KH_2_PO_4_, 42 mM Na_2_HPO_4_, 8.5 mM NaCl, 18.7 mM NH_4_Cl,1 mM MgSO_4_) and recorded for 30 s after 30 s acclimatisation on a Leica S9i microscope at 26 fps. All strains were recorded until day 8, except GMC101 which did not survive until this age. Recorded videos were analysed using the wrMTrck plugin of ImageJ to quantify body bends per second in an unbiased way [33]. The experiment was carried out in a blinded manner. For each condition around 30 animals were analysed.

### NIAD-4 staining

An age-synchronised population was grown at 20 °C until reaching adulthood, after which the temperature was switched to 25 °C until day 3 of adulthood. Animals were washed in M9 buffer and incubated in 1 µM NIAD-4 (Cayman Chemicals) dye solution in M9 buffer for 4 h. They were washed with M9 and destained on a seeded NGM plate for 16 h after which they were imaged as described below.

### Microscopy and FRAP

Nematodes were transferred to 2.5 % agarose pads on microscopy slides containing 5 mM tetramisole hydrochloride (Sigma Aldrich) as an anaesthetic agent. Immobilised animals were imaged by microscopy within 1 h.

Widefield imaging was performed on a Leica DMi8 inverted microscope with N Plan 10x/0.25 PH1 or HC PL FL L 20x/0.40 Ph1 air objectives.

Confocal microscopy was performed on a Nikon Eclipse Ti2 body equipped with an X-Light V3 spinning disk (Crest Optics) and Nikon Plan Apo 60xA/1.40 oil DIC H oil objective. The excitation and the emission wavelengths were set to 470 nm and 525/50nm respectively to visualise GFP. NIAD-4 was imaged using 470 nm and 605/70 nm excitation and emission wavelengths, respectively.

For FRAP, age-synchronised day 3 animals were immobilised on microscopy slides as stated above. A Nikon Eclipse Ti microscope body equipped with CSU-X1-A1 spinning disk (Yokogawa) and Nikon Plan Apo VC 60x /1.40 oil objective was used. Imaging was done at an excitation wavelength of 488 nm using 10 % laser power and the region of interest was bleached with 100 % laser power for 10 repetitions. Regions of interest for photobleaching were imaged for 0.8 s before bleaching and 24 s after bleaching. Signal intensities for each time point were measured by ImageJ and normalised to t_0_ signal intensity.

### Hexanediol treatment

For hexanediol treatment, nematodes were grown until day 4 of adulthood at 20°C. Animals were immobilised on microscope slides in 20 µL thermo-reversible hydrogel (CyGel^TM^, BioStatus) containing 10 % 1,6-hexanediol (Sigma Aldrich) and 10 mM tetramisole hydrochloride (Sigma Aldrich). Embedded animals were incubated for 1 h at room temperature, after which confocal imaging was performed as described above.

### Western blot and fractionation

Age-synchronised nematodes were collected in M9 buffer on day 1 of adulthood and snap-frozen in liquid nitrogen. 5X Sample buffer (5 % SDS, 50 % glycerol, 0.1 % Bromophenol blue, 250 mM Tris-HCl pH 6.8 and 5 % β-Mercaptoethanol) was added and samples were boiled at 90 °C for 10 minutes. Proteins were separated on a 10 % SDS-PAGE gel and transferred to a PVDF membrane using the Power Blotter System (Invitrogen). The following primary antibody was used in this study: mouse anti-GFP (JL8, Takara). Goat anti-mouse (A32730, Invitrogen) was used as secondary antibody and blots were scanned on an Odyssey DLx Imager (Li-Cor).

For the fractionation assay, ca. 5000 adult animals per sample were harvested and snap-frozen in liquid nitrogen. Pellets were resuspended in RIPA lysis buffer (ThermoFisher Scientific) containing protease inhibitor cocktail (Roche) and lysed using the TissueLyser II (Qiagen) for 5 minutes at a frequency of 30 s^-1^. The supernatant was fractionated into SDS-soluble and insoluble fractions by centrifugation at 20,800 g for 30 minutes at 4 °C. The pellet was washed with RIPA buffer once and dissolved in urea buffer (8 M urea, 2 % SDS, 50 mM DTT, 50 mM Tris-HCl pH 8). All fractions were run on SDS-PAGE followed by western blot as described above.

### Immunoprecipitation and negative staining TEM

For the immunoprecipitation assay, ca. 5000 adult worms per sample were harvested and snap-frozen in liquid nitrogen. Pellets were resuspended in native lysis buffer (50 mM Tris-HCl pH 7.4, 5 mM MgCl_2_, 0.5 % Triton-X 100, 0.2 mM PMSF, 1 µg/mL Leupeptin, 1 µg/mL Pepstatin A) containing protease inhibitor cocktail (Roche) and lysed using the TissueLyser II (Qiagen) for 5 minutes at a frequency of 30 s^-1^. The lysate was applied to GFP-Trap Agarose beads (Chromotek) equilibrated in the same buffer. The purification was done according to the manufacturer’s guide.

For negative staining, 4 μL of the purified sample was spotted onto 200 mesh formvar/carbon grids, which were previously glow discharged for 15 s at 10 mA. Samples were incubated for 1 minute, washed 2 times with water, and finally negatively stained with 2 % uranyl acetate for 1 minute. The grids were imaged on a Tecnai 20 electron microscope (FEI Company) operated at 200 kV with a LaB6 filament and FEI Eagle CCD camera. Images were acquired using FEI TEM Imaging & Analysis (TIA) software.

### Statistical analysis

For statistical analysis, GraphPad Prism 9 was used. Statistical analysis was performed using two-way ANOVA analysis with Dunnet multiple comparison test for the motility assays, and using one-way ANOVA with Tukey test for the western blot results. Data are presented as the mean ± standard deviation. p ≤ 0.05 was used as the threshold for considering the data statistically significant.

## Acknowledgement

We thank Jorieke Tiggelaar and Maithili Joshi for performing *C. elegans* micro-injections, Barend Elenbaas for providing the synthetic IAPP sample, the Biology Imaging Centre and Martin Haase at Physical and Colloidal Chemistry (Utrecht University) for the use of the confocal microscopes, and Menno Bergmeijer and the Utrecht University Electron Microscopy Centre for support and access to TEM. We also thank Antoinette Killian and the members of the Sinnige lab for valuable discussions.

## Author contributions

Mehmet Akdag: conceptualisation, investigation, formal analysis, visualisation, writing – original draft. Vera van Schijndel: investigation, visualisation, writing – review & editing. Tessa Sinnige: conceptualisation, supervision, writing – original draft.

## References

[1] International Diabetes Federation, IDF Diabetes Atlas 2021 _ IDF Diabetes Atlas, IDF Official Website. (2021).

[2] B. Åkesson, G. Panagiotidis, P. Westermark, I. Lundquist, Islet amyloid polypeptide inhibits glucagon release and exerts a dual action on insulin release from isolated islets, Regul Pept. 111 (2003) 55–60. 10.1016/S0167-0115(02)00252-5.

[3] P. Westermark, A. Andersson, G.T. Westermark, Islet Amyloid Polypeptide, Islet Amyloid, and Diabetes Mellitus, Physiol Rev. 91 (2011) 795–826. 10.1152/physrev.00042.2009.

[4] S. Asthana, B. Mallick, A.T. Alexandrescu, S. Jha, IAPP in type II diabetes: Basic research on structure, molecular interactions, and disease mechanisms suggests potential intervention strategies, Biochimica et Biophysica Acta (BBA) - Biomembranes. 1860 (2018) 1765–1782. 10.1016/j.bbamem.2018.02.020.

[5] C. Huang, C. Lin, L. Haataja, T. Gurlo, A.E. Butler, R.A. Rizza, P.C. Butler, High Expression Rates of Human Islet Amyloid Polypeptide Induce Endoplasmic Reticulum Stress–Mediated β-Cell Apoptosis, a Characteristic of Humans With Type 2 but Not Type 1 Diabetes, Diabetes. 56 (2007) 2016–2027. 10.2337/db07-0197.

[6] A.E. Butler, J. Janson, S. Bonner-Weir, R. Ritzel, R.A. Rizza, P.C. Butler, β-Cell Deficit and Increased β-Cell Apoptosis in Humans With Type 2 Diabetes, Diabetes. 52 (2003) 102–110. 10.2337/diabetes.52.1.102.

[7] D.C. Rodriguez Camargo, S. Chia, J. Menzies, B. Mannini, G. Meisl, M. Lundqvist, C. Pohl, K. Bernfur, V. Lattanzi, J. Habchi, S.I. Cohen, T.P.J. Knowles, M. Vendruscolo, S. Linse, Surface-Catalyzed Secondary Nucleation Dominates the Generation of Toxic IAPP Aggregates, Front Mol Biosci. 8 (2021). 10.3389/fmolb.2021.757425.

[8] B.O.W. Elenbaas, L. Khemtemourian, J.A. Killian, T. Sinnige, Membrane-Catalyzed Aggregation of Islet Amyloid Polypeptide Is Dominated by Secondary Nucleation, Biochemistry. 61 (2022). 10.1021/acs.biochem.2c00184.

[9] Y. Xu, R. Maya-Martinez, N. Guthertz, G.R. Heath, I.W. Manfield, A.L. Breeze, F. Sobott, R. Foster, S.E. Radford, Tuning the rate of aggregation of hIAPP into amyloid using small-molecule modulators of assembly, Nat Commun. 13 (2022) 1–15. 10.1038/s41467-022-28660-7.

[10] M.F.M. Engel, L. Khemtémourian, C.C. Kleijer, H.J.D. Meeldijk, J. Jacobs, A.J. Verkleij, B. De Kruijff, J.A. Killian, J.W.M. Höppener, Membrane damage by human islet amyloid polypeptide through fibril growth at the membrane, Proc Natl Acad Sci U S A. (2008). 10.1073/pnas.0708354105.

[11] P. Cao, A. Abedini, H. Wang, L.-H. Tu, X. Zhang, A.M. Schmidt, D.P. Raleigh, Islet amyloid polypeptide toxicity and membrane interactions, Proceedings of the National Academy of Sciences. 110 (2013) 19279–19284. 10.1073/pnas.1305517110.

[12] B.O.W. Elenbaas, S.M. Kremsreiter, L. Khemtemourian, J.A. Killian, T. Sinnige, Fibril elongation by human islet amyloid polypeptide is the main event linking aggregation to membrane damage, BBA Advances. 3 (2023) 100083. 10.1016/j.bbadva.2023.100083.

[13] J.W.M. Höppener, C. Oosterwijk, M.G. Nieuwenhuis, G. Posthuma, J.H.H. Thijssen, Th.M. Vroom, B. Ahrén, C.J.M. Lips, Extensive islet amyloid formation is induced by development of Type II diabetes mellitus and contributes to its progression: pathogenesis of diabetes in a mouse model, Diabetologia. 42 (1999) 427–434. 10.1007/s001250051175.

[14] H.J. Hiddinga, S. Sakagashira, M. Ishigame, P. Madde, T. Sanke, K. Nanjo, Y.C. Kudva, J.J. Lee, J. Van Deursen, N.L. Eberhardt, Expression of wild-type and mutant S20G hIAPP in physiologic knock-in mouse models fails to induce islet amyloid formation, but induces mild glucose intolerance, J Diabetes Investig. 3 (2012) 93–198. 10.1111/j.2040-1124.2011.

[15] C.D. Link, Expression of human β-amyloid peptide in transgenic Caenorhabditis elegans, Proc Natl Acad Sci U S A. 92 (1995) 9368–9372. 10.1073/pnas.92.20.9368.

[16] G. McColl, B.R. Roberts, T.L. Pukala, V.B. Kenche, C.M. Roberts, C.D. Link, T.M. Ryan, C.L. Masters, K.J. Barnham, A.I. Bush, R.A. Cherny, Utility of an improved model of amyloid-beta (Aβ1-42) toxicity in Caenorhabditis elegans for drug screening for Alzheimer’s disease, Mol Neurodegener. (2012). 10.1186/1750-1326-7-57.

[17] H.R. Brignull, F.E. Moore, S.J. Tang, R.I. Morimoto, Polyglutamine Proteins at the Pathogenic Threshold Display Neuron-Specific Aggregation in a Pan-Neuronal *Caenorhabditis elegans* Model, The Journal of Neuroscience. 26 (2006) 7597–7606. 10.1523/JNEUROSCI.0990-06.2006.

[18] T.J. van Ham, K.L. Thijssen, R. Breitling, R.M.W.W. Hofstra, R.H.A.A. Plasterk, E.A.A.A. Nollen, C. elegans Model Identifies Genetic Modifiers of α-Synuclein Inclusion Formation During Aging, PLoS Genet. 4 (2008) e1000027. 10.1371/journal.pgen.1000027.

[19] J.F. Morley, H.R. Brignull, J.J. Weyers, R.I. Morimoto, The threshold for polyglutamine-expansion protein aggregation and cellular toxicity is dynamic and influenced by aging in Caenorhabditis elegans., Proc Natl Acad Sci U S A. 99 (2002) 10417–22. 10.1073/pnas.152161099.

[20] M. Brehme, C. Voisine, T. Rolland, S. Wachi, J.H. Soper, Y. Zhu, K. Orton, A. Villella, D. Garza, M. Vidal, H. Ge, R.I. Morimoto, A Chaperome Subnetwork Safeguards Proteostasis in Aging and Neurodegenerative Disease, Cell Rep. 9 (2014) 1135–1150. 10.1016/j.celrep.2014.09.042.

[21] B. Calamini, M.C. Silva, F. Madoux, D.M. Hutt, S. Khanna, M. a Chalfant, S.A. Saldanha, P. Hodder, B.D. Tait, D. Garza, W.E. Balch, R.I. Morimoto, Small-molecule proteostasis regulators for protein conformational diseases, Nat Chem Biol. 8 (2012) 185–196. 10.1038/nchembio.763.

[22] J. Habchi, P. Arosio, M. Perni, A.R. Costa, M. Yagi-Utsumi, P. Joshi, S.K.R. Chia, S.I.A. Cohen, M.B.D. Müller, S. Linse, E.A.A. Nollen, C.M. Dobson, T.P.J. Knowles, M. Vendruscolo, An anti-cancer drug suppresses the primary nucleation reaction that initiates the formation of toxic Aβ aggregates associated with Alzheimer’s disease, Sci Adv. 2 (2016) e1501244. 10.1126/sciadv.1501244.

[23] R.F. Laine, T. Sinnige, K.Y. Ma, A.J. Haack, C. Poudel, P. Gaida, N. Curry, M. Perni, E.A.A.A. Nollen, C.M. Dobson, M. Vendruscolo, G.S.K. Schierle, C.F. Kaminski, G.S. Kaminski Schierle, C.F. Kaminski, Fast fluorescence lifetime imaging reveals the aggregation processes of α-synuclein and polyglutamine in aging Caenorhabditis elegans, ACS Chem Biol. 414714 (2019) 1628–1636. 10.1021/acschembio.9b00354.

[24] Y. Aldras, S. Singh, K. Bode, D.C. Bhowmick, A. Jeremic, D.M. O’Halloran, An inducible model of human amylin overexpression reveals diverse transcriptional changes, Neurosci Lett. 704 (2019) 212–219. 10.1016/j.neulet.2019.04.016.

[25] P.C. Rosas, G.M. Nagaraja, P. Kaur, A. Panossian, G. Wickman, L.R. Garcia, F.A. Al-Khamis, A.A.A. Asea, Hsp72 (HSPA1A) prevents human islet amyloid polypeptide aggregation and Toxicity: A new approach for type 2 diabetes treatment, PLoS One. (2016). 10.1371/journal.pone.0149409.

[26] N.C. Kegulian, S. Sankhagowit, M. Apostolidou, S.A. Jayasinghe, N. Malmstadt, P.C. Butler, R. Langen, Membrane curvature-sensing and curvature-inducing activity of islet amyloid polypeptide and its implications for membrane disruption, Journal of Biological Chemistry. 290 (2015). 10.1074/jbc.M115.659797.

[27] T. Gurlo, S. Ryazantsev, C.J. Huang, M.W. Yeh, H.A. Reber, O.J. Hines, T.D. O’Brien, C.G. Glabe, P.C. Butler, Evidence for proteotoxicity in β cells in type 2 diabetes: Toxic islet amyloid polypeptide oligomers form intracellularly in the secretory pathway, American Journal of Pathology. 176 (2010). 10.2353/ajpath.2010.090532.

[28] A. Ahier, C.Y. Dai, A. Tweedie, A. Bezawork-Geleta, I. Kirmes, S. Zuryn, Affinity purification of cell-specific mitochondria from whole animals resolves patterns of genetic mosaicism, Nat Cell Biol. 20 (2018). 10.1038/s41556-017-0023-x.

[29] E.E. Nesterov, J. Skoch, B.T. Hyman, W.E. Klunk, B.J. Bacskai, T.M. Swager, In vivo optical imaging of amyloid aggregates in brain: Design of fluorescent markers, Angewandte Chemie - International Edition. 44 (2005). 10.1002/anie.200500845.

[30] J.R. Brender, J. Krishnamoorthy, M.F.M. Sciacca, S. Vivekanandan, L. D’Urso, J. Chen, C. La Rosa, A. Ramamoorthy, Probing the Sources of the Apparent Irreproducibility of Amyloid Formation: Drastic Changes in Kinetics and a Switch in Mechanism Due to Micellelike Oligomer Formation at Critical Concentrations of IAPP, J Phys Chem B. 119 (2015) 2886–2896. 10.1021/jp511758w.

[31] C. Gallrein, M. Iburg, T. Michelberger, A. Kocak, D. Puchkov, F. Liu, S.M.A. Mariscal, T. Nayak, G.S.K. Schierle, J. Kirstein, Novel Amyloid-beta pathology C. elegans model reveals distinct neurons as seeds of pathogenicity, Prog Neurobiol. 198 (2020) 107229. 10.1016/j.buildenv.2020.107229.

[32] T. Sinnige, P. Ciryam, S. Casford, C.M. Dobson, M. De Bono, M. Vendruscolo, Expression of the amyloid-β peptide in a single pair of C. elegans sensory neurons modulates the associated behavioural response, PLoS One. (2019). 10.1371/journal.pone.0217746

[33] C.I. Nussbaum-Krammer, M.F. Neto, R.M. Brielmann, J.S. Pedersen, R.I. Morimoto, Investigating the Spreading and Toxicity of Prion-like Proteins Using the Metazoan Model Organism C. elegans, Journal of Visualized Experiments. (2015). 10.3791/52321.

