## Supplementary Information for "Islet amyloid polypeptide tagged with green fluorescent protein localises to mitochondria and forms filamentous aggregates in *Caenorhabditis elegans*"

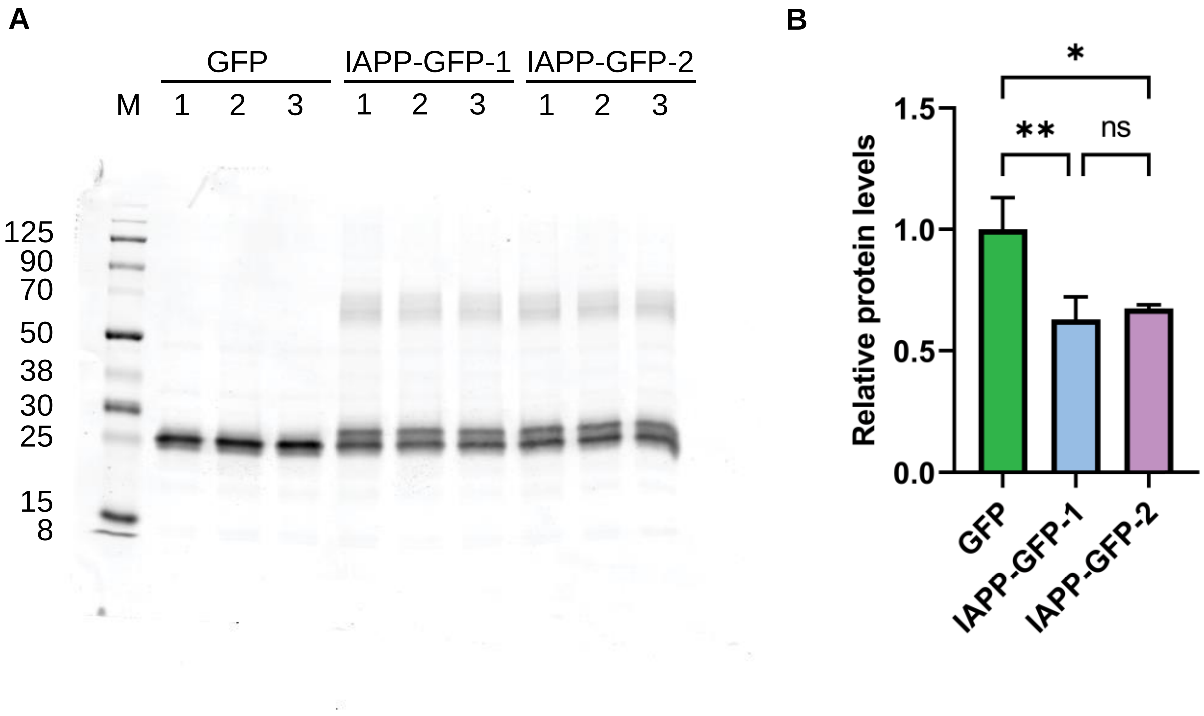


**Figure S1:** IAPP-GFP strains show similar protein expression levels. **(A)** IAPP-GFP protein expression validated by GFP western blot. Worm lysates were separated on SDS-PAGE and membranes were blotted with an anti-GFP antibody. **(B)** Quantification of the western blot. IAPP-GFP runs as a double band, the lower one corresponding to the size of GFP which is presumably a cleavage product. Both bands were included in the quantification. One-way ANOVA analysis with Tukey multiple comparison test was used for analysis. (* $p < 0.05$; ** $p < 0.01$)

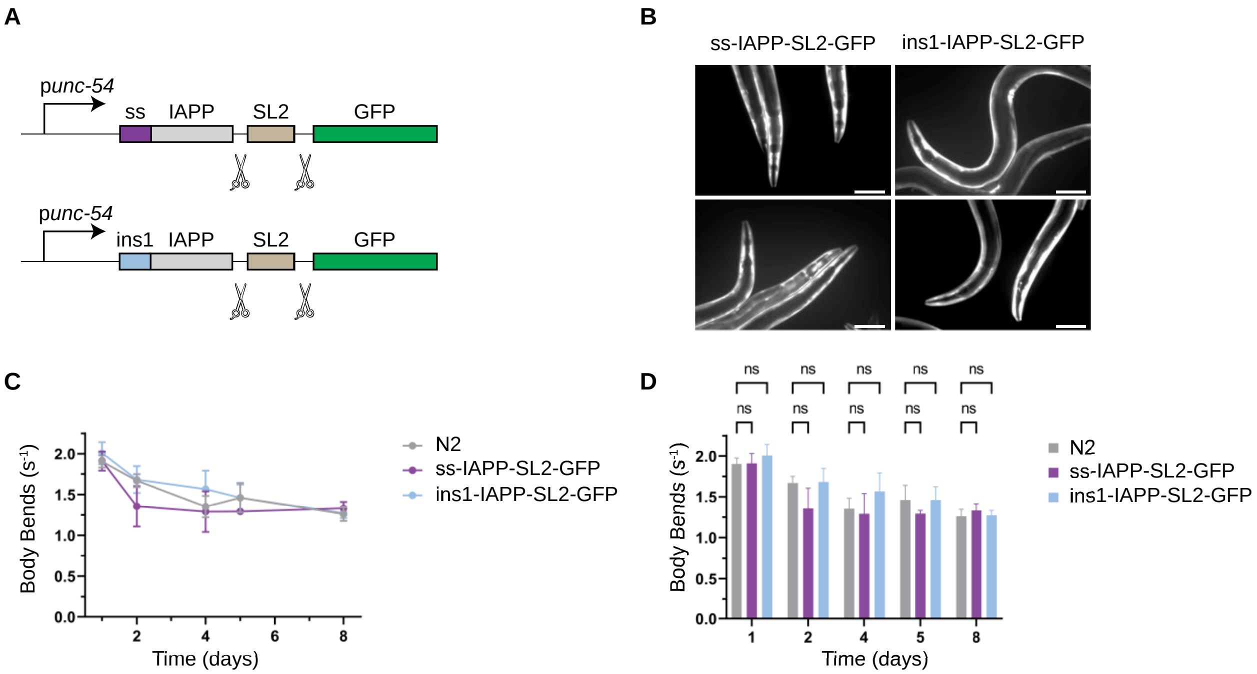


**Figure S2**: Construction of body wall muscle cell specific *C. elegans* IAPP-SL2-GFP models. **(A)** Schematic design of the constructs used to generate ss-IAPP-SL2-GFP and ins1-IAPP-SL2-GFP strains. **(B)** Fluorescence microscopy images validating the expression of GFP in the body wall muscle cells. (Scale bar 100 µm) **(C)** Motility assay to monitor protein toxicity. Wild-type (N2) strain was used as the control group. Data collection and analysis were done in a blinded manner and at least 30 worms were used for each data point. **(D)** Quantification of the motility assay. Two-way ANOVA analysis with Dunnet multiple comparison test was employed for statistical analysis. (* $p < 0.05$; ** $p < 0.01$)

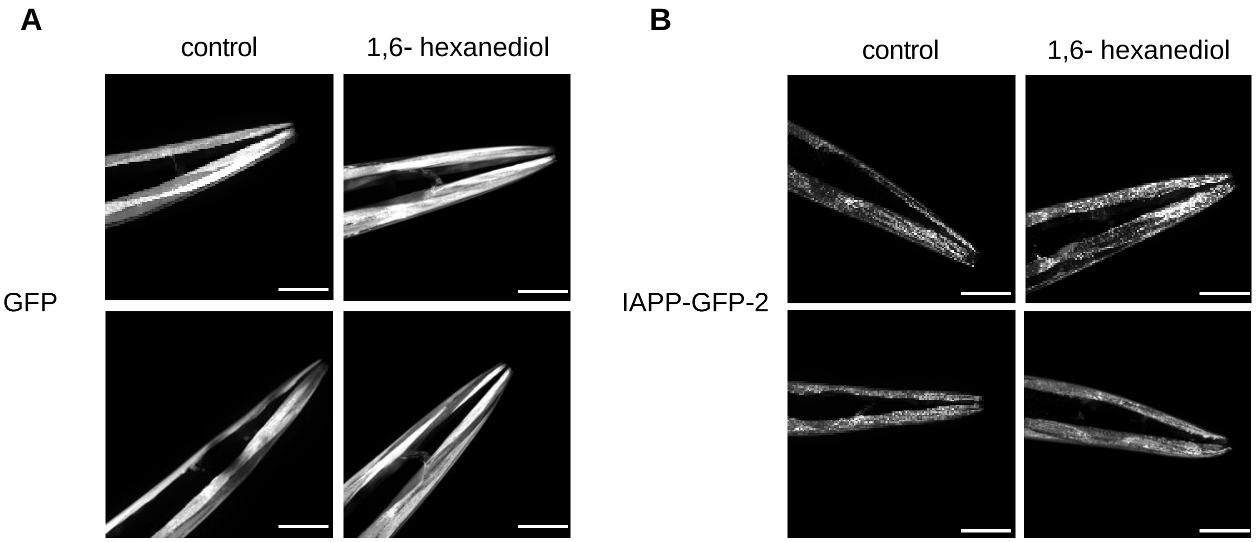


**Figure S3:** IAPP-GFP distribution does not change by 1,6-hexanediol treatment. 1,6-hexanediol treated **(A)** GFP and **(B)** IAPP-GFP-2 strains showed similar distribution as mock treated control groups. (Scale bar 50 µm)

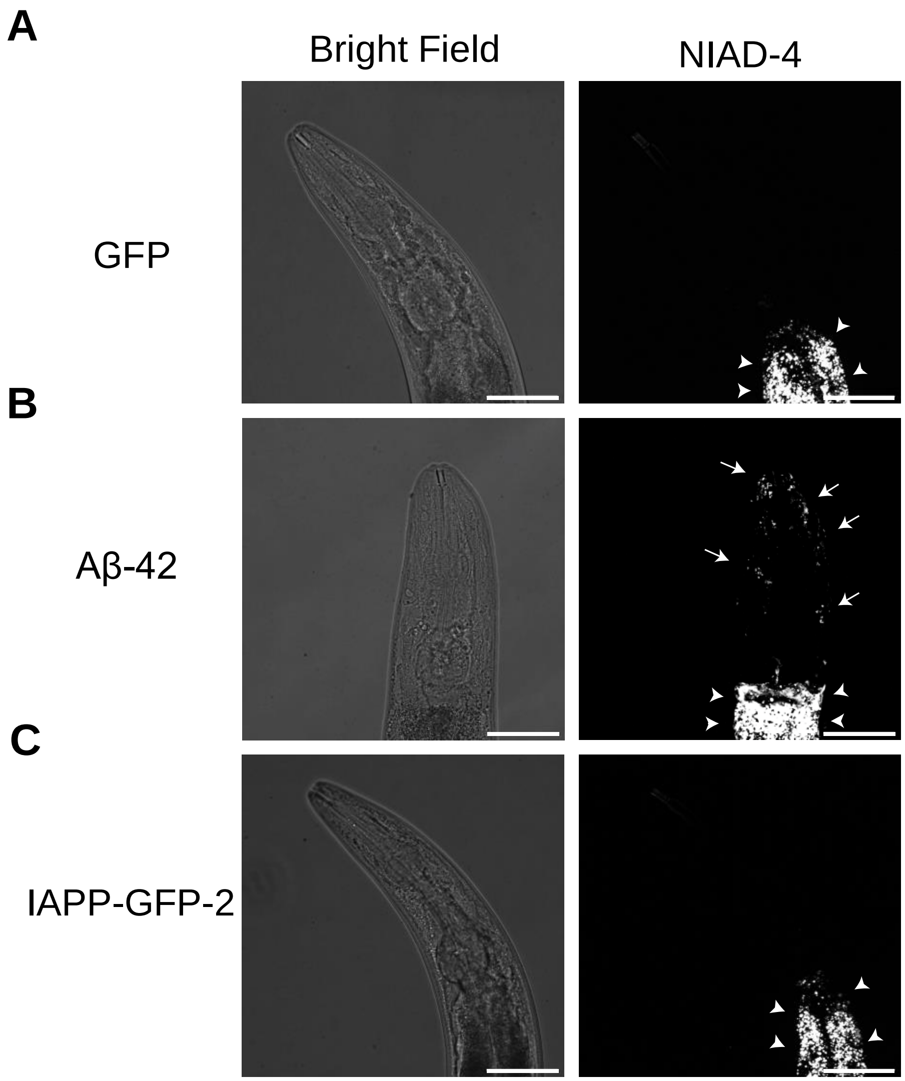


**Figure S4:** Representative images of NIAD-4 stained amyloid deposits in **(A)** GFP, **(B)** Aβ-42 and **(C)** IAPP-GFP-2 strains. Only Aβ-42 showed NIAD-4 specific amyloid deposits (arrows). The remaining bright signal corresponds to unspecific staining of the intestine (arrow heads). (Scale bar 50 µm)

**Table S1:** Plasmids used in this study

| Plasmid | Description |
|---|---|
| pDEST R4-R3 Vector II (Invitrogen) | Ampicillin resistant general cloning vector |
| pMA05 | Body wall muscle cell specific ss-IAPP-SL2-GFP expression plasmid |
| pMA06 | Body wall muscle cell specific ins1-IAPP-SL2-GFP expression plasmid |
| pMA08 | Body wall muscle cell specific IAPP-GFP expression plasmid |
| pMA09 | Body wall muscle specific GFP expression plasmid |

**Table S2:** Oligonucleotides used to generate IAPP-GFP construct (pMA08)

| Primer | Sequence |
|---|---|
| *unc-54*p fwd | 5'- aacatatccagtcactatgGCTGCAGTGAGTATTTTCGG- 3' |
| *unc-54* rev | 5'- catttttctgaaaagcctgctacgtCTCGCTTCTTTCAAATGGTT- 3' |
| IAPP fwd | 5'- acgtagcaggcttttcagaaaaATGAAATGCAACACTGCCACATG- 3' |
| IAPP rev | 5'-gcccactttgtacaagaaagctgggtcATATGTATTGGATCCCACGTTGG- 3' |
| GFP fwd | 5'-gacccagctttcttgtacaaagtgggcATGAGTAAAGGAGAAGAACTTTTC- 3' |
| *unc-54* 3'UTR rev | 5'-gagaaaataccgcatcaggcGGCCGACTAGTAGGAAACAG - 3' |

**Table S3:** Oligonucleotides used to generate IAPP-SL2-GFP constructs (pMA05 and pMA06)

| **Primer** | **Sequence** |
|---|---|
| SL2 fwd | 5’ -tgggatccaatacatattaaGCTGTCTCATCCTACTTTCACC- 3’ |
| SL2 rev | 5’ -ctttactcattttttctaccggtacaGCAGTTTC- 3’ |
| GFP fwd | 5’ -ggtagaaaaaATGAGTAAAGGAGAAGAACTTTTCACTG- 3’ |
| *unc-54* 3’UTR rev | 5’ -gagaaaataccgcatcaggcGGCCGACTAGTAGGAAACAG- 3’ |
| ins1-IAPP oligo sequence | 5’-ttttcagaaaaATGTACTGGTTTCGTCAAGTTTACAGACCCTCGTTCTTCTTTGGCTTTCTCGCGATCCTTCTCCTCTCGTCGCCGACGCCTTCAGACGCAAAATGCAACACTGCCACATGTGCAACGCAGCGCCTGGCAAATTTTTTAGTTCATTCCAGCAACAACTTTGGTGCCATTCTCTCATCTACCAACGTGGGATCCAATACATATtaa |
| ss-IAPP oligo sequence | 5’- ttttcagaaaaATGCATAAGGTTTTGCTGGCACTGTTCTTTATCTTTCTGGCACCAGCAGGTACCGACGCGAAATGCAACACTGCCACATGTGCAACGCAGCGCCTGGCAAATTTTTTAGTTCATTCCAGCAACAACTTTGGTGCCATTCTCTCATCTACCAACGTGGGATCCAATACATATtaa |
